# Transection of the ventral hippocampal commissure impairs spatial reference but not contextual or spatial working memory

**DOI:** 10.1101/437491

**Authors:** Jake T. Jordan, Yi Tong, Carolyn L. Pytte

**Affiliations:** Biology Department, The Graduate Center, City University of New York (CUNY), New York, NY 11016; CUNY Neuroscience Collaborative, The Graduate Center, CUNY, New York, NY 11016; Psychology Department, Queens College, CUNY, Flushing, NY 11367; Psychology Department, The Graduate Center, CUNY, New York, NY 11016

## Abstract

Plasticity is a neural phenomenon in which experience induces long-lasting changes to neuronal circuits and is at the center of most neurobiological theories of learning and memory. However, too much plasticity is maladaptive and must be balanced with substrate stability. Area CA3 of the hippocampus is lateralized with the left hemisphere dominant in plasticity and the right specialized for stability. Left and right CA3 project bilaterally to CA1; however, it is not known whether this downstream merging of lateralized plasticity and stability is functional. We hypothesized that interhemispheric integration of input from these pathways is essential for integrating spatial memory stored in the left CA3 with spatial working memory facilitated by the right CA3. To test this, we severed interhemispheric connections between the left and right hippocampi in mice and assessed learning and memory. Despite damage to this major hippocampal fiber tract, hippocampus-dependent spatial working memory and short- and long-term memory were both spared. However, tasks that required the integration of information retrieved from memory with ongoing spatial working memory and navigation were impaired. We propose that one function of interhemispheric communication in the mouse hippocampus is to integrate lateralized processing of plastic and stable circuits to facilitate memory-guided spatial navigation.

## Introduction

The hippocampus is essential for spatial memory and navigation and is a site of robust synaptic plasticity. Synaptic plasticity manifests as enduring experience-driven changes to the strength of communication between neurons and has long been a central tenant of neurobiological theories of memory [1-3]. Many studies have investigated the mechanisms by which neural circuits balance plasticity with stability [4]; however, it is not clear whether plasticity and stability are required to perform specific neural computations.

Long-term potentiation (LTP) is a form of synaptic plasticity that has been extensively studied since its discovery at synapses between hippocampal areas CA3 and CA1 almost five decades ago [5]. Unlike traditional electrophysiological stimulation, optogenetic tools allow for targeted stimulation of specific populations of axons in brain regions, such as the CA1, that receive interleaved inputs from multiple sources [6]. Optogenetic studies have revealed that LTP is left-lateralized in the mouse hippocampus such that left CA3 sends highly plastic projections to both left and right CA1 while right CA3 sends highly stable projections to both left and right CA1 [6,7]. This lateralized specialization underscores the need for understanding the mouse hippocampus as a bilateral structure [7-10]. In particular, it is not yet known whether or how lateralized plastic and stable circuits play a role in hippocampus-dependent cognition.

The hippocampus sends direct interhemispheric projections along the entire dorsoventral hippocampus via a structure called the ventral hippocampal commissure (VHC). Interhemispheric synthesis of lateralized processing is a key component of hypotheses describing the function of the bilateral hippocampus [8,10,11]. Recently, we have proposed that the plastic left CA3 acquires, stores and retrieves spatial and contextual information, discretizing such information into representations of particular places. Further, the stable right CA3 may be part of a spatial working memory system that computes and continuously updates routes during spatial navigation. The VHC enables this lateralized processing to be projected to bilateral CA1, integrating left-lateralized memory retrieval with right-lateralized spatial working memory and navigation to facilitate memory-guided spatial navigation (Fig. 1A; [10]). Spatial memory tasks that require both retrieval of learned places and navigation through those places would require interhemispheric communication and would be impaired by loss of VHC function (Fig. 1B). Tasks that require retrieval of learned places or contexts but that do not require spatial navigation would be unimpaired by VHC damage. Likewise, tasks that require spatial working memory only but not retrieval of previously learned places or contexts would not require the VHC. Other hypotheses of the bilateral hippocampus have proposed roles of the VHC in interhemispheric transfer of spatial memory engrams [11] and in facilitating short-term memory by increasing bilateral processing power [7]. Despite the critical role of interhemispheric communication in all of these hypotheses, no study has yet examined the role of interhippocampal pathways in learning and memory in mice.

**Fig. 1.**
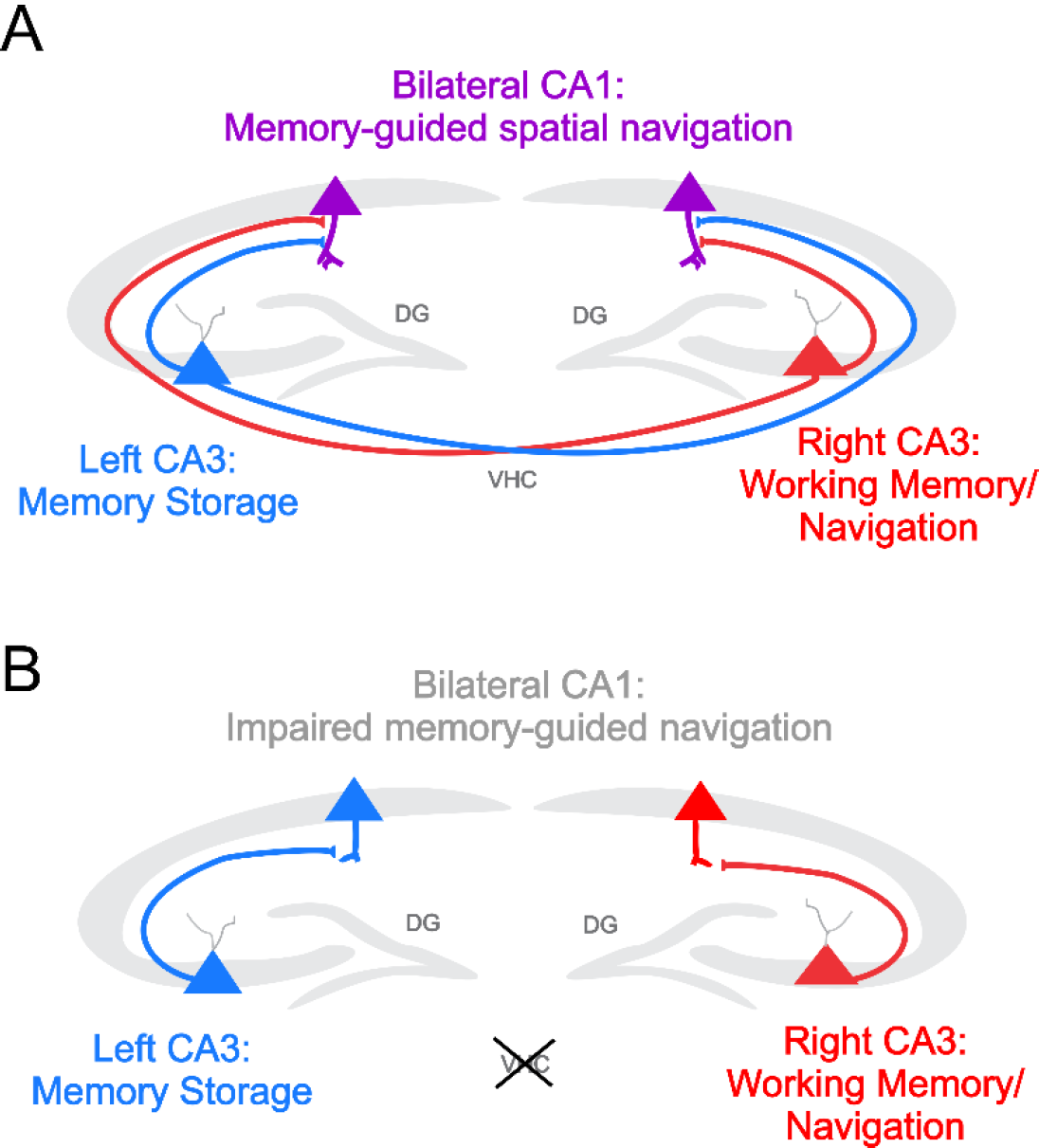
Hypothesized effects of VHC transection. **A**, Our hypothesis is that plastic left CA3 projections (blue) facilitate memory storage and stable right CA3 projections (red) facilitate spatial working memory. Bilateral projections from left and right CA3 would integrate spatial information stored in left CA3 with navigation computations in right CA3 in bilateral CA1, thus facilitating spatial navigation through remembered environments (purple). **B**, Loss of the ventral hippocampal commissure results in a loss of integration such that memory-guided spatial navigation would be impaired, while memory retrieval and spatial working memory would be unaffected.

To test these hypotheses and to learn more about the role of interhemispheric communication in hippocampal cognitive function, we developed a surgical approach for severing interhemispheric fibers in mice. We tested these mice on a series of hippocampus-dependent tasks selected to test key predictions of proposed models of the bilateral hippocampus. We found that memory retrieval in the absence of navigation was unimpaired, in contrast to predictions of an interhemispheric transfer of hippocampal engrams [11]. Further, there was no consistent impairment of short-term or working memory, suggesting that interhemispheric communication does not enable short-term and working memory [7]. Instead, we found a consistent pattern of impairments on tasks that required both memory retrieval and spatial navigation, consistent with our hypothesis of lateralized memory and navigation [10].

## Results

We severed inter-hemispheric pathways by sectioning the VHC in C57BL6J mice (S1 Fig., Online Methods). The VHC is a pathway that contains inter-hemispheric projections along the entire dorsoventral extent of both hippocampi. Sham split-brain surgeries consisted of a posterior to anterior incision in the cortex parallel to the midline, immediately left or right of the superior sagittal sinus, and dorsal to the corpus callosum (CC), leaving all interhemispheric fibers intact. In cc mice, CC fibers dorsal to the VHC were sectioned in addition to overlying cortex with no damage to the VHC. In vhc+cc mice, the VHC was sectioned in addition to the overlying CC and cortex (S1B Fig. and Online Methods). We then compared hippocampus-dependent memory using a series of behavioral paradigms across treatment groups.

We sought to determine if retrieval of past experiences would be spared in split-brain mice, which we predicted is a left-lateralized process that would not require interhemispheric communication (Fig. 1B). Contextual fear memory is a widely-used hippocampus-dependent task in which mice are allowed to explore a chamber before receiving a mild foot shock. The strength of the memory for the context is measured by re-exposing the conditioned mouse to that chamber at a later time and measuring the time spent freezing, a naturally occurring fear response. We trained mice on a contextual fear conditioning protocol known to be particularly sensitive to hippocampal manipulations as lesions of the dorsal hippocampus and even ablation of adult hippocampal neurogenesis produced anterograde amnesia of contextual fear memory when only a single shock was given during conditioning [12-15]. We conducted retrieval sessions one (short-term) and 24 hours (long-term) after conditioning in which the animal was returned to the context in which they had received a mild foot shock (Fig. 2A). There was no apparent difference in time spent freezing across treatment groups in short- or long-term retrieval sessions occurring one (Fig. 2B) and 24 hours (Fig. 2C) after conditioning. We then sought to determine whether split-brain surgery would also spare spatial working memory, which we predicted is a right-lateralized process that would not require interhemispheric communication (Fig. 1B). We performed a spontaneous alternation task in mice freely exploring a Y-Maze apparatus in a single trial (Fig. 2D). In this task, rates of alternation between arms (as opposed to the revisiting of recently explored arms) is assessed. Bilateral lesions of the dorsal hippocampus have been shown to reduce the percentage of exploration events considered alternations without reducing the total number of exploration events [16]. We found no effect of surgery on percent alternation in this task (Fig. 2E). Further, there was no effect of surgery on the total number of exploration events (S2 Fig.), indicating that a lack of differences in percent alternation was not masked by differences in amount of exploration. These data indicate that inter-hemispheric communication between the left and right hippocampi was not needed for short- and long-term contextual memory retrieval or for spatial working memory.

**Fig. 2.**
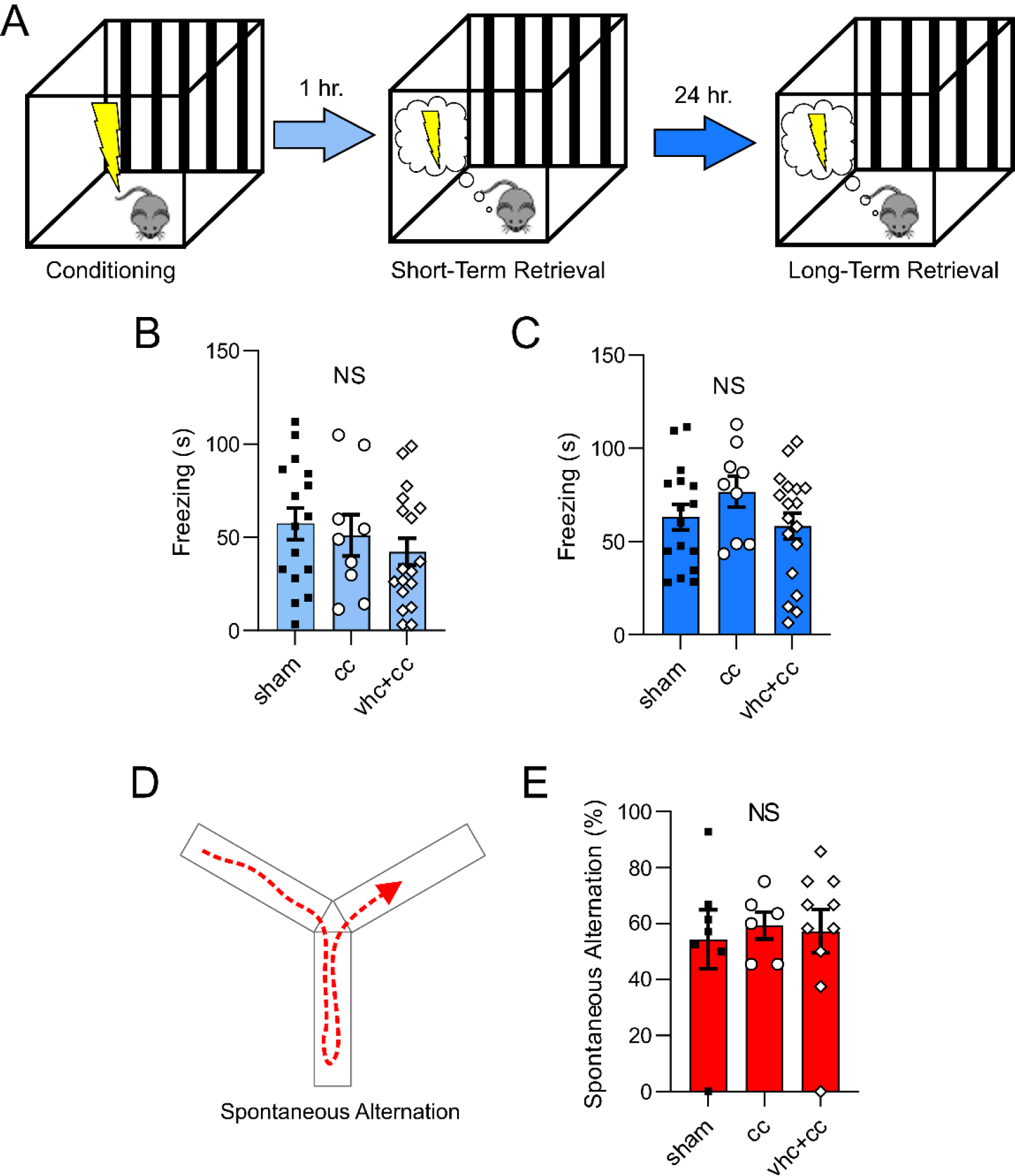
vhc+cc transection spares contextual fear and spatial working memory. **A**, Hippocampus-dependent contextual fear memory paradigm. Mice were place in a behavioral chamber where they received a single mild foot-shock during a conditioning session. Mice were re-exposed to the chamber one and 24 hours later to asses short- and long-term contextual fear memory (n = 16 sham, 9 cc, 18 vhc+cc). **B**, Time spent freezing during context re-exposure one hour after conditioning. There was no effect of surgical treatment on freezing (F(2,40) = 0.920; *p* = 0.401). **C**, Time spent freezing during context re-exposure 24 hours after conditioning. There was no effect of surgical treatment on freezing (F(2,40) = 1.316; *p* = 0.280). **D**, Mice explored a Y-Maze apparatus to assess the percentage of navigation events classified as spontaneous alternation (n = 7 sham, 6 cc, 10 vhc+cc). **E**, There was no effect of surgical treatment on percent alternation (F(2,20) = 0.076; *p* = 0.927). Error bars represent SEM; dots indicate individual mice; NS indicates no significance.

We then sought to determine if split-brain surgery affected behavior during a task that integrated memory retrieval with spatial working memory. We tested mice on a short-term memory version of the Y-Maze task that is equally impaired following inhibition of either the left or right CA3 (Fig. 3A). Mice explored two arms of a three-arm Y-Maze for two minutes with the third arm blocked off. Mice were removed from the Y-Maze for one minute and then re-exposed for a two-minute retrieval trial in which all arms were available for exploration. Mice typically exhibit a preference for novelty and although they tend to explore the entire maze during the retrieval trial, they typically spend the most time in the novel arm. We first asked whether there was any difference in preference for the novel arm at the beginning of the trial. Across groups, there was no difference in the proportion of mice that explored the novel arm first (6 of 9 sham mice, 8 of 9 cc mice, 8 of 10 vhc+cc mice; *Χ*^2^ = 1.339, *p* = 0.512). We then compared the latency to explore the novel arm across treatments. Although there was a trend towards a difference among groups (Fig. 3B), this did not appear to be driven by effects of vhc+cc surgery but rather by a decreased latency in the cc group. The implication of this potential difference is not clear. However, preference for the novel arm at the beginning of the retrieval trial did not appear to be affected by vhc+cc transection, indicating intact short-term retrieval of the familiar arm (t(16) = 1.239, x, p = 0.233). We then measured preference for the novel arm over the familiar arm throughout the retrieval trial by comparing the total time spent in the novel over the familiar arm. As expected, sham and cc mice exhibited a strong preference for the novel arm over the familiar arm during the retrieval trial. Vhc+cc mice showed no such preference (Fig. 3C) indicating that although they initially demonstrated a preference for the novel arm that was comparable to the other groups, indicating intact short-term memory retrieval, this preference diminished as exploration of the maze took place.

**Fig. 3.**
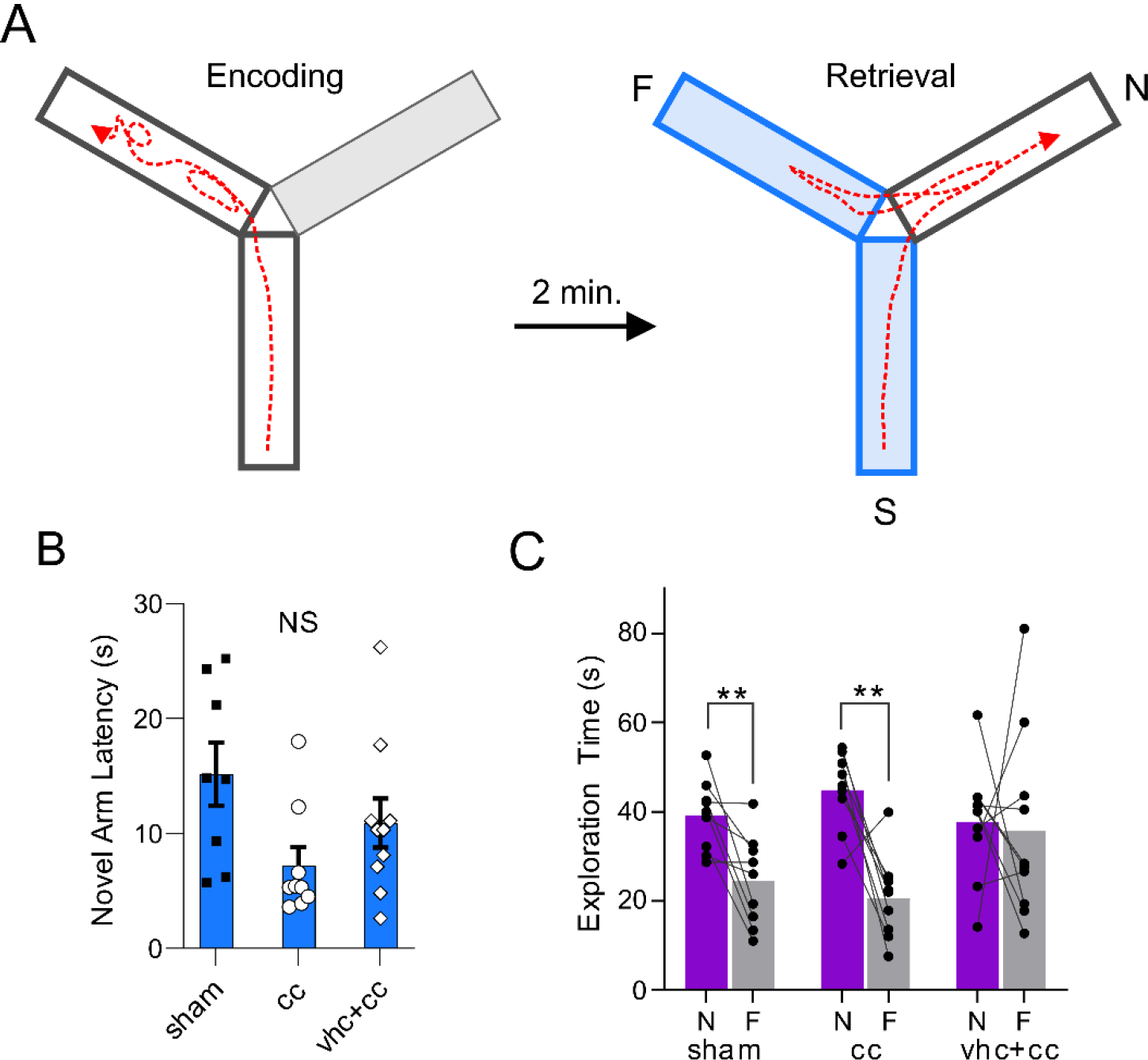
vhc+cc transection spares immediate recognition of spatial novelty but impairs preference for spatial novelty with exploration. **A**, Y-Maze short-term spatial memory paradigm. During an encoding trial, mice were allowed to explore two arms of a three-arm Y-Maze. Blocked arm is shown in green. Following a 1-minute intertrial interval, mice were returned to the apparatus with all arms open. S indicates the start arm; F indicates the familiar arm that the mouse had access to during the encoding trial; N indicates the novel arm that was blocked off during encoding (n = 9 sham, 9 cc, 10 vhc+cc). **B**, Latency to explore the novel arm was not significantly different across treatment groups (F(2,24) = 3.162, *p* = 0.060). **C**, Time spent in the novel and familiar arms during the retrieval trial. Sham and cc mice spent significantly more time exploring the novel arm than the familiar arm (sham: t(8) = 3.502, *p* = 0.008; cc: t(8) = 4.641, *p* = 0.002). vhc+cc mice exhibited no such preference (t(9) = 0.205; *p* = 0.842). Error bars represent SEM; dots indicate individual mice; \*\**p* < 0.01, NS indicates no significance.

The Morris Water Maze (MWM) is a widely-used long-term spatial learning and memory paradigm that requires both retrieval of previously learned information as well as spatial navigation and thus tests our prediction that this task would be impaired following loss of VHC function. We sought to determine if interhemispheric communication was required for memory-guided navigation in the long-term MWM task. Mice were trained on a spatial version of the MWM in which the escape platform was hidden below the surface of the water and could be found using distal spatial cues (Fig. 4A). In rats, both the left and right hippocampi are needed for the spatial MWM [17], though they offer distinct, complementary contributions to this task [11]. Spatial acquisition was assessed by measuring the average escape latency across four trials per day, each day over five days of training. Although each group demonstrated a decrease in escape latencies throughout training, vhc+cc mice had longer escape latencies than those of both sham and cc mice (Fig. 4B). Thus, all treatment groups exhibited spatial learning, however vhc+cc mice showed impaired spatial learning. This occurred despite intact learning on a non-spatial version of the MWM that is hippocampus-independent (S3 Fig.). Twenty-four hours after the end of spatial MWM acquisition training, a 60-second probe trial was conducted in which the escape platform was removed and time spent searching in the target quadrant was measured. Sham and cc mice searched selectively, exhibiting a preference for the target quadrant above chance levels. Vhc+cc mice showed no memory for platform location, showing chance levels of search in the target quadrant, indicating that vhc+cc mice did not appear to distinguish between the target quadrant and others (Fig. 4C). Thus, loss of VHC function in mice impairs long-term spatial learning and memory on the MWM.

**Fig. 4.**
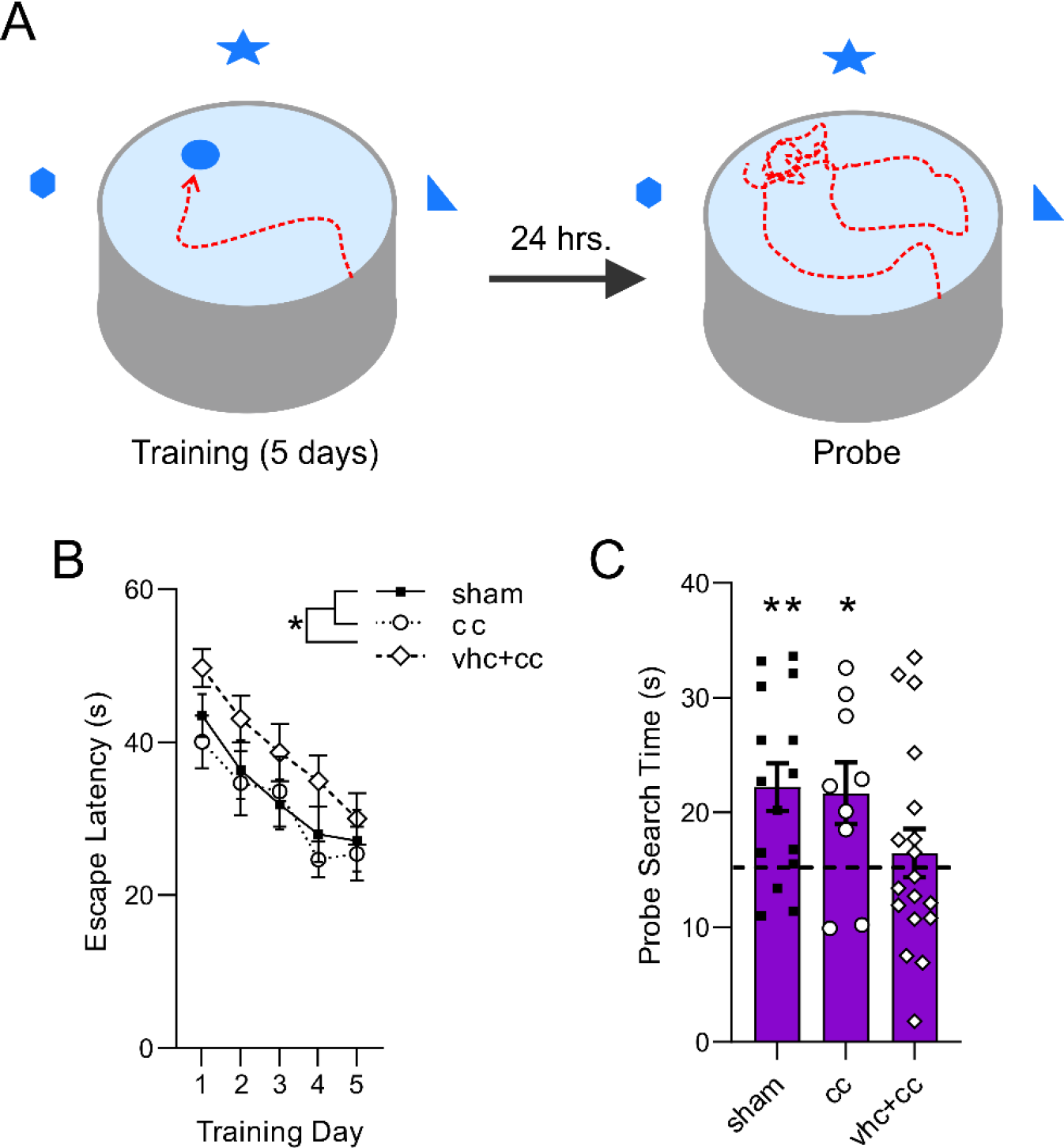
vhc+cc transection impairs long-term spatial memory acquisition and retrieval. **A**, MWM spatial learning and memory paradigm. Mice were trained to locate a hidden escape platform (purple circle) by its relation to extra-maze spatial cues. After 5 days of training, mice were given a single probe trial with the escape platform removed (target quadrant shown in purple, n = 15 sham, 9 cc, 18 vhc+cc). **B**, Average escape latencies on each day of training. We found a main effect of training day (F(4,156) = 12.75, *p* < 0.0001) and of treatment group (F(2,39) = 3.575, *p* = 0.032) on escape latency during acquisition, with no interaction between these factors (F(8,156) = 0.208, *p* = 0.989). Post hoc analyses showed that vhc+cc mice were significantly slower than sham (vhc+cc vs. sham: p = 0.031) and cc mice (vhc+cc vs. cc: *p* = 0.006). Sham mice did not differ from cc mice during acquisition (*p* = 0.772). **C**, Time spent searching in the target quadrant for each treatment group on the probe trial. Sham and cc mice searched selectively, exhibiting a preference for the target quadrant above chance levels (sham: t(14) = 3.510, *p* = 0.004; cc: t(8) = 2.480, *p* = 0.038). vhc+cc mice did not appear to distinguish between the target quadrant and others (t(17) = 0.694; *p* = 0.497). Error bars represent SEM; dots indicate individual mice; \**p* < 0.05, \*\**p* < 0.01.

To rule out potential effects of split-brain surgery on vision, locomotion, motivation, or non-hippocampus-dependent procedural learning, mice were trained on a version of the MWM in which the platform was visible throughout the duration of each trial. There was no effect of surgical treatment on escape latencies throughout training (S3A Fig.). As an additional test of non-specific surgical effects, we compared shortest latencies at the end of training to determine whether fastest trials differed across groups. There were no differences across treatments (S3B Fig.). Earlier work found that split-brain mice with monocular deprivation showed increased avoidance behavior, possibly influencing the interpretation of deficits seen in behavioral paradigms [18]. To determine whether split-brain surgery affected avoidance behavior, we exposed mice to a single session of exploration of an elevated plus maze. There was no difference in open arm avoidance behavior across treatment groups measured as time spent in open arms or as bouts of open arm exploration (S4 Fig.).

## Discussion

Transection of the VHC and CC, but not of the CC alone, impaired performance of certain hippocampus-dependent behavioral tasks while sparing others. Specifically, loss of VHC function had no effect on short- or long-term contextual fear memory or on Y-Maze spatial working memory (Fig. 2). After exposure to two arms of a Y-Maze, initial preference for the novel arm following a short delay was unaffected by loss of VHC function indicating intact short-term spatial memory. However, preference for the novel arm over the course of the entire retrieval trial was greatly impaired by loss of VHC function suggesting that integration of memory retrieval with working memory as mice explored the maze requires interhemispheric communication via the VHC. Finally, loss of VHC function impaired long-term spatial learning and memory on the MWM. These impairments were not likely due to aberrant effects of split-brain surgery on vision, locomotion, motivation, hippocampus-independent learning or avoidance behavior. We believe it is unlikely that cellular remodeling following ventral hippocampal commissure transection contributed to the impairments seen in our spatial memory experiments as the contextual fear conditioning protocol we used reveals functional manipulations of hippocampal circuitry [12-15]. Further, although our vhc+cc surgery necessarily damaged structures overlying the VHC, such as the fimbria, fornix, and septum (S1B Fig.), contextual fear memory has been shown to be impaired following lesions to the fimbria and fornix [19], and contextual fear expression is suppressed during inhibition of the septum [20]. Because we found no effect of either the cc or vhc+cc surgery on contextual fear encoding or expression, we suggest that any damage done to these structures as a result of surgery was not sufficient to impair this behavior. Taken together, our data indicate inter-hippocampal communication is not required for the ability to acquire, store, and retrieve new hippocampal memories nor for the ability to retain spatial information in working memory. However, it is required for performance of tasks that specifically integrate retrieved spatial memories with spatial working memory and navigation.

Left and right CA3 present a provocative opportunity to understand the functions of plasticity and stability in cognition. Left CA3 projections to bilateral CA1 are highly plastic in mice [6,7] and are required for long-term spatial memory acquisition [7,21], consistent with a long-theorized role of plasticity in memory formation [2]. In contrast to left CA3, right CA3 projections to bilateral CA1 are highly stable in mice [6,7] and both left and right CA3 are needed for short-term spatial memory [7]. Thus, it was suggested that interhemispheric convergence of CA3 input in bilateral CA1 facilitates short-term memory [7]. Over time, long-term memory no longer requires right CA3 as synaptic consolidation occurs at highly plastic left CA3 synapses with bilateral CA1 [7]. Loss of VHC function would therefore impair short-term spatial memory while long-term memory would be spared. However, our data do not support this hypothesis as short-term contextual fear memory and recognition of the novel arm on the Y-Maze short-term spatial memory task were unaffected by loss of VHC function. Further, the requirement of inter-hippocampal communication on the MWM long-term spatial memory task suggests that the left hippocampus alone cannot perform the necessary processing required for this task. The requirement of inter-hippocampal communication for the spatial MWM also conflicts with the conclusions of a study which suggested that spatial memory is right-dominant in the hippocampus [18]. This work did not inactivate the hippocampus, but rather studied the effects of monocular deprivation on Barnes Maze spatial memory retrieval and therefore, their results may be explained in terms of lateralized visuospatial processing.

Our data are consistent with our model of the bilateral hippocampus in which left and right CA3 have distinct but complementary functions [10]. In addition to the studies in mice, discussed above, we also based this hypothesis on the findings in humans and rats that the right hippocampus is specialized for navigation through learned environments [11,22]. Although the hemispheric differences in synaptic plasticity that have been reported in mice using optogenetic tools have not been reported in humans or rats, we propose that lateralized hippocampal function is largely homologous across these three species. Across these three species, the left hippocampus appears to be essential for acquiring new spatial memories [7,11,22], which again, is consistent with a long-theorized role of plasticity in memory [2] and is largely agreed upon in hypotheses regarding hippocampal lateralization [7,8,10,11]. The function of the synaptically stable right CA3 is not yet clear. The requirement of the right CA3 in short-term memory tasks in mice and the requirement of the right hippocampus for navigation in humans and rats may be explained by a common property of right CA3 function. We have proposed that the right CA3 facilitates navigation by specializing in the computation of route-related metric information such as distance and direction, as reported in human neuroimaging studies [23,24]. This information is held in a spatial working memory system during navigation. In this case, any task that requires spatial working memory and/or navigation through an environment will require a contribution from right CA3.

In summary, we have demonstrated a role for inter-hippocampal communication in hippocampus-dependent tasks that require both hippocampus-dependent memory retrieval and spatial navigation. We propose that hippocampal plasticity and stability have complementary functions in memory and route computation, respectively, and can perform these functions in isolation of each other. Integration of lateralized processing by plastic and stable hippocampal circuits is required for integrating memory retrieval with route computation to facilitate memory-guided spatial navigation.

## Acknowledgments

We thank Steven Jett for help with myelin staining; Jeff Beeler, and Dan McCloskey for input on experimental design; Joshua Brumberg, and Tiago Gonçalves for comments on the manuscript. This work was supported by the Alfred P Sloan Foundation (76332-28) and PSC-CUNY TRADA-43-687 to CLP and CUNY Doctoral Student Research Grant and Mina Rees Dissertation Fellowship to JTJ.

## Supplemental Methods

### Animals

We used adult male and female C57/BL6J mice (Jackson Labs) that had been bred in-house for 2-5 generations. All mice were adults aged 2.5-7 months at the time of surgery. Mice were housed on a 12 h light/dark cycle, with all behavioral sessions occurring during the light phase. All procedures were approved by the Queens College Institutional Animal Care and Use Committee (protocol # 177).

### Experimental Design

The ventral hippocampal commissure contains axons connecting the left and right hippocampi. The term ventral hippocampal commissure (vhc) distinguishes this structure from the dorsal hippocampal commissure, which connects extra-hippocampal cortical areas in the left and right hemisphere and should not be confused with the ventral/dorsal distinction of the hippocampus itself^25^. Fibers in the vhc originate and terminate throughout the entire dorsal-ventral extent of the hippocampus. We performed “complete” or “partial” split-brain surgeries. A complete split-brain surgery consisted of transection of both the vhc and the overlying corpus callosum (cc), as the vhc cannot be accessed without transecting the cc. Partial split-brain surgery consisted of transection of only the cc located over the vhc to control for possible contributions of the cc to the behavioral assays, and hippocampus-dependent memory more specifically^26^. Sham split brain surgeries consisted of sectioning cortex overlying the cc as occurred in both the treatment groups. We refer to mice receiving complete split-brain surgery as vhc+cc (n = 11; 4 females, 7 males), mice receiving partial split-brain surgery as cc (n = 9; 5 females, 4 males), and sham split-brain mice as sham (n = 9; 4 females, 5 males). There were no sex differences in any behavioral measure, therefore males and females were combined for all analyses. These mice were trained and tested in: 1) short-term spatial memory Y-Maze, 2) Morris Water Maze, 3) short- and long-term contextual fear memory, and 4) spatial working memory Y-Maze.

Twenty-eight additional male mice were used in an experiment to quantify immediate early gene expression in sham and vhc+cc mice (Npas4 expression data not shown in this manuscript). Of these, 14 mice were trained and tested in the Morris Water Maze (n = 7 sham, 7 vhc+cc) and 14 mice were used in the short- and long-term contextual fear experiment (n = 7 sham, 7 vhc+cc). As the behaviors of these mice did not differ from those of the first cohort, we combined the behavioral data of both cohorts for analysis. Sample sizes were corrected following histological confirmation of the surgeries and varied by behavioral test, see individual sections in Results.

### Surgery

In order to sever inter-hemispheric pathways, we modified a method developed by Schalomon & Wahlsten^27^. We used an L-shaped, sharpened piece of tungsten wire (0.25 mm in diameter) as a knife. The body temperature of the mice was monitored and maintained with a heating pad. Mice were anesthetized with ketamine/xylazine (intraperitoneal, 90-120 mg/kg and 5-10 mg/kg, respectively) and further anesthetized for one minute in an isoflurane chamber (4.5% isoflurane in oxygen). Mice were placed in a stereotaxic apparatus, receiving a constant flow of 1.5% isoflurane and oxygen (1.5 L/minute) and were given an injection of bupivacaine (1.25-2 mg/kg) under the scalp for local analgesia. An opening was made by drilling two adjacent 1 mm-wide holes into the skull to access the brain. To avoid the superior sagittal sinus, openings in the skull were made 0.5 mm lateral to the midline and the side of surgery for each animal was randomly chosen (±0.5 ML, −0.8 AP from bregma for the first hole, ±0.5 ML, −1.6 AP from bregma for the second hole). To sever both the VHC and CC, the short, sharpened end of the L-knife was placed on the surface of the brain along the medial side of the hole and was then slowly lowered 3.5 mm with the joint of the L along the posterior edge of the hole and the short arm of the blade facing anteriorly. Once lowered, the knife was translated anteriorly so that the knife moved posterior to anterior to “hook” the VHC and CC fibers. The knife was then raised 2.5 mm until the short arm reached 1 mm below the underside of the skull. The knife was then translated back and raised out of the hole. To sever the CC only, we performed the same procedure as vhc+cc transection, however the knife was only lowered 2.2 mm. For sham surgeries, we used the same procedure but the knife was lowered 1.0 mm. Mice were administered buprenorphine following surgery (0.1 mg/kg, subcutaneous). Transected and sham mice were indistinguishable by blind observation of their behavior in the home cage.

### Behavior

Behavioral testing began approximately four weeks after surgery. Mice were habituated to handling for one day by the experimenter. On testing days, mice were transported to a designated behavior room and were allowed to acclimate for 20-30 minutes before the start of the task. All behavior was scored by an experimenter blind to surgical condition and sex using the Stopwatch+ program.

Short-term spatial memory was measured using a Y-Maze task established to be equally sensitive to inactivation of either the left or the right hippocampus^7^. The Y-maze was constructed of clear acrylic and had three arms (height: 20 cm; length: 30 cm; width: 8 cm) 120 degrees apart. The room contained many spatial cues including light fixtures and furniture. In addition, a painting and a movie poster were placed on the walls in line with the axes of the familiar and novel arms. The experimenter stood along the axis of the start arm during each trial. Each arm was marked with a black line at the entrance to determine whether the mouse was in the arm or not. Mice were considered to be in an arm if all four paws were across the entrance line. The paradigm consisted of a 2-minute encoding trial during which one arm was blocked off, followed by a 1-minute intertrial interval, then a 2-minute retrieval session. The start arm remained the same in both the encoding and retrieval trial, while the exposed arm during the encoding trial was considered the familiar arm and the blocked arm was considered the novel arm. At the start of the encoding trial, mice were placed facing outward in the start arm and were allowed to explore the start and familiar arms for two minutes, beginning when the mouse left the start arm. Mice were removed from the Y-Maze and placed back into their home cage for one minute during the intertrial interval. While mice were in the home cage, the block was removed to expose the novel arm. To remove odor cues, the apparatus was wiped with 70% ethanol, rotated 120 degrees, and then wiped dry with a paper towel before the retrieval trial. After the intertrial interval, the mice were again placed facing out in the start arm and were allowed to explore the entire maze for two minutes. At the end of the retrieval session, mice were placed back in their home cages and the maze wiped and dried before the next animal was placed in the maze. Y-Maze short-term spatial memory was scored as the latency to explore the novel arm as well as a comparison between the times spent in the novel and familiar arms during the retrieval paradigm. One vhc+cc mouse was removed from analysis for failure to leave the start arm during the retrieval trial and therefore had scores of zero for both the novel and familiar arms.

Long-term spatial learning and memory was measured using the Morris Water Maze (MWM), following the protocol of Vorhees & Williams^28^. The pool was 110 cm in diameter and was filled with opaque water colored with non-toxic white paint maintained at a temperature of 25.3°C ± 0.5°C. The escape platform was white and was submerged approximately 0.5 cm under the surface of the water. The platform remained in the same location throughout training. Salient room cues were visible from the surface of the pool and included colored and patterned posters, lighting, and furniture. Training consisted of four trials per day for five days with the starting location varying on each trial. Intertrial intervals were 30 seconds, during which the mice remained on the platform before starting the next trial. Mice that did not reach the platform within 60 seconds were placed onto the platform. Twenty-four hours after the final training, a 60-second probe trial was given during which the escape platform was removed.

To measure spatial learning, latencies to the escape platform were recorded for each training trial and were averaged across trials for each mouse on each of the five training days. For the probe trials, spatial memory was scored by time spent in the target quadrant versus chance levels of performance^11^. One sham mouse was removed from analysis for exhibiting signs of hypothermia after a training session.

Hippocampus-independent learning was assessed using a version of the MWM in which the escape platform was visible, as described by Vorhees & Williams^28^. Mice were tested in a different room but used the same pool as in the spatial paradigm. However, the platform was above water level and a red disk was placed on top to contrast the platform with the white pool and water. Training consisted of three trials per day over five days. Mice were placed in the water facing the wall on the opposite side of the pool from the target platform. After finding the platform, mice were left for 15 seconds before being moved to the home cage for the intertrial interval and the platform was moved to a new spatial location. The intertrial interval had no set time and ended when the platform was moved and the water settled (approximately 30 seconds). If mice did not find the platform within 60 seconds, they were placed onto it by the experimenter and remained for 30 seconds.

Spatial working memory was tested using a Y-Maze spatial working memory task sensitive to dorsal hippocampal lesions^16^. The Y-Maze apparatus used was the same as used in the short-term spatial memory task described above. Each arm was demarcated by a black line that distinguished the arm and formed a triangle in the center of the maze. A mouse was considered to have left an arm when all four paws crossed this line heading into the center of the maze and was considered to have entered an arm when all four paws crossed the line heading into the arm. The maze was positioned on a table or desk at least one meter from extra-maze spatial cues. Spatial cues consisted of posters on the walls, furniture, experimental equipment and the experimenter. The maze was wiped down with 70% ethanol and dried before each trial. Trials were two minutes each and began when the mouse was placed facing outward at the end of a randomly chosen arm. Each navigation event (when a mouse left an arm and then entered an arm) over the course of the two-minute trial was scored as one of the following: 1) a ‘same arm return’ was counted when a mouse left an arm and then returned to that same arm 2) a ‘revisit’ was counted when a mouse left an arm and returned to the arm it had explored previously, and 3) an ‘alternation’ was counted when a mouse left an arm and returned to an arm it had not previously been in. The first choice in the session was not recorded; the first choice after a same arm return was not recorded. After two minutes, the mouse was removed from the maze and placed back into its home cage. Spatial working memory was scored by the percent of navigation events scored as alternation^16^.

Short- and long-term contextual fear memory was tested using a one-shock conditioning paradigm that is sensitive to hippocampal manipulations^12-15^. Conditioning took place in a fear conditioning chamber housed in a sound-attenuating cubicle (Med Associates). Before each session, the chamber was wiped down with 70% ethanol and dried. On the first day, mice were placed in the chamber and allowed to explore freely for three minutes, then given a mild foot-shock (2 s, 0.75 mA), and removed 15 seconds later (total conditioning session time = three minutes 17 seconds). Two 3-minute retrieval sessions occurred one and 24 hours after conditioning, a protocol previously used to dissociate molecular contributions to short-and long-term contextual fear memory^29^. Fear expression was scored as time spent freezing during each minute of the three-minute retrieval trial (absence of all movement, except breathing).

Avoidance behavior was measured using the elevated plus maze, a paradigm sensitive to both dorsal and ventral hippocampal manipulations^25,30^. The elevated plus apparatus consisted of four arms (30.5 cm long, 6.4 cm wide), two of which were enclosed on three sides with walls (20.3 cm high). Mice were placed in the center of the apparatus and allowed to explore freely for 5 minutes and were then returned to their home cage. The apparatus was wiped with 70% ethanol and dried both before and after each trial. Avoidance behavior was scored as time with all four paws on an open arm as well as number of entries into the open arms. One male vhc+cc mice was removed from analysis as it fell off the apparatus during the session.

### Myelin Staining and Surgical Verification

Surgeries were verified 4-6 months after surgery. Mice were euthanized with 0.3 mL of euthasol. Brains were extracted and post-fixed in 4% paraformaldehyde. Brains were then cryoprotected in 30% sucrose before cryosectioning coronally at 60 µm. To assess transection to the cc and vhc, we stained sections with luxol blue and cresyl violet. Slides were dried overnight at 37°C. Histology began with a de-fat step in which sections were serially dehydrated in ethanol and then placed in xylene (twice for five minutes each). Sections were then rehydrated and submersed in 70% ethanol for one hour at room temperature. Sections were then incubated in a 0.1% luxol blue solution in 95% ethanol overnight at 56°C. Myelin was differentiated via rinses in deionized water, followed by 0.05% lithium carbonated in deionized water, followed by 70% ethanol (two minutes each; differentiation was repeated as necessary). Sections were then stained in 0.1% cresyl violet in deionized water, serially dehydrated, cleared in xylene and coverslipped using Krystalon.

### Statistical Analysis

To assess spatial working memory, we compared the percent of alternations as well as number of events across treatment groups using one-way ANOVAs for each measure. To assess short-term memory during the Y-Maze retrieval trial, we planned comparisons to determine whether each treatment group exhibited a preference for the novel arm over the familiar arm, but not whether treatment groups differed in exploration times. Therefore, we compared time spent in the novel arm to time spent in the familiar arm within each group using Student’s paired t-tests. We also assessed novelty to explore the novel arm with a one-way ANOVA. One outlier was removed from this analysis (female sham) as its latency was greater than two standard deviations greater than the mean. To assess cued and spatial learning on the MWM, we performed two-way mixed model ANOVAs (treatment × training day, with training day as the repeating factor) on escape latency, followed by post hoc tests (Tukey’s HSD). To assess spatial memory on the 60-second MWM probe trial, we used one-sample t-tests within each treatment group, comparing time spent in the target quadrant to chance performance (15 seconds)^11^. We used a one-way ANOVA to determine whether minimum escape latencies differed among groups on the final day of visible MWM training. To assess short- and long-term contextual fear memory, we performed a one-way ANOVA across treatment groups on time spent freezing during each of the two retrieval trials. To assess avoidance behavior during the elevated plus maze test, we performed a one-way ANOVA on time spent in the open arms across all groups.

---

**Supplementary Figure 1.**
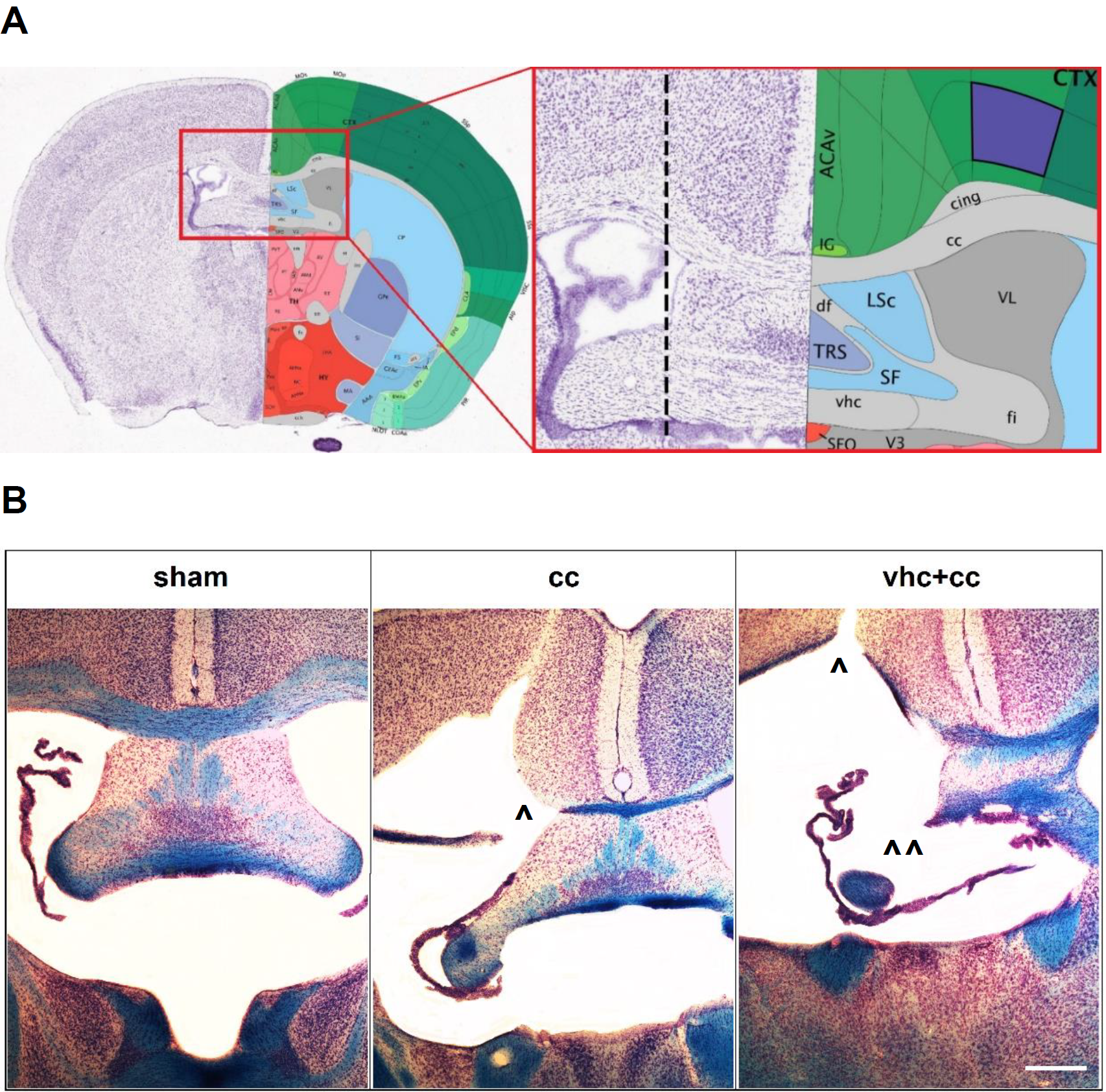
**A**, Anatomical illustration of a bilateral coronal section showing site of surgical treatments. Dotted line indicates path of surgical knife. ACAv, ventral anterior cingulate cortex; cing, cingulum bundle; cc, corpus callosum; CTX, cortex; df, dorsal fornix; fi, fimbria; IG, induseum griseum; LSc, TRS, and SF are all nuclei of the septum; SFO, subfornical organ; V3, third ventricle; vhc, ventral hippocampal commissure; VL, lateral ventricle. Image credit: Allen Institute, © 2004 Allen Institute for Brain Science. Allen Mouse Brain Atlas, available from http://mouse.brain-map.org/ **B**, Surgical treatment groups. Myelin is stained blue (luxol fast blue) with cresyl violet counterstaining. B, Sham surgery consisted of lowering the surgical knife into the cortex, leaving the corpus callosum (cc) and ventral hippocampal commissure (vhc) intact; cc surgery consisted of transection of the cc, sparing the vhc; vhc+cc surgery consisted of transection of both the ventral hippocampal commissure and overlying corpus callosum. ^ transection of the cc, ^ ^ transection of the vhc, scale bar = 500 μm.

**Supplementary Figure 2.**
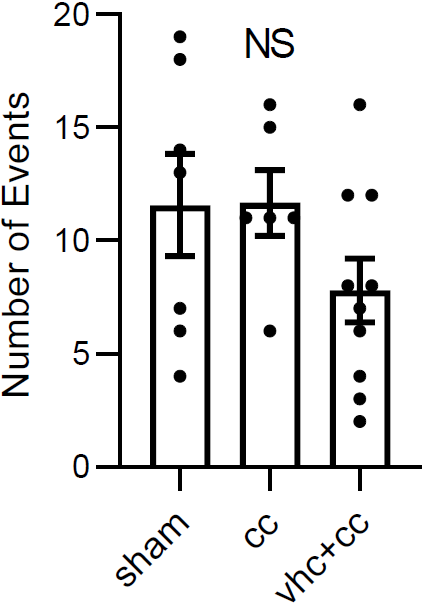
Total number of navigation events in the Y-Maze working memory task was not affected by surgical treatment (F(2,20) = 1.810, *p* = 0.189).

**Supplementary Figure 3.**
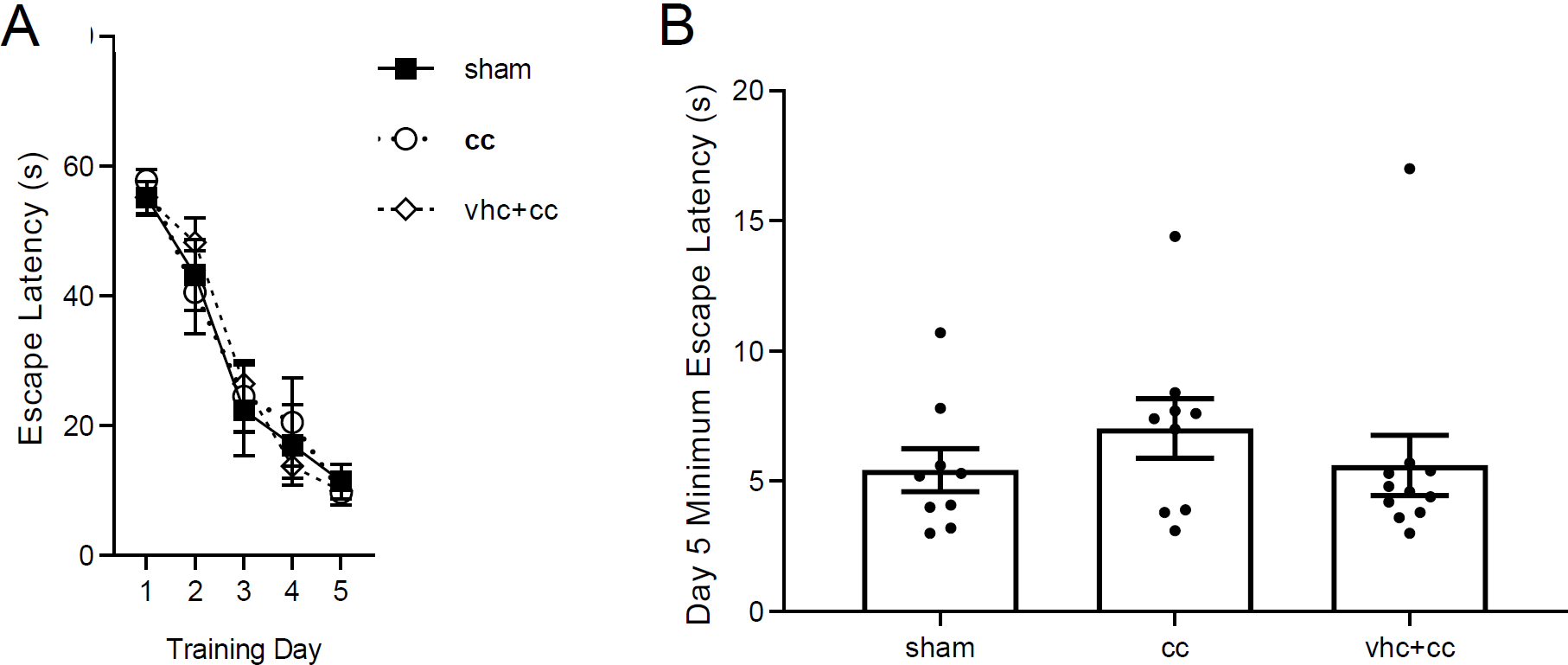
Hippocampus-independent MWM acquisition with a visible platform. **A**, We found a main effect of training day on escape latency (F(4,112) = 98.1, *p* < 0.0001), no effect of treatment (F(2,26) = 0.024, *p* = 0.976), and no interaction between these factors (F(8,112) = 0.766, *p* = 0.6332). **B**, Minimum escape latencies on the final day of training did not differ among treatments (F(2,26) = 0.632, *p* = 0.539). Sham: n = 11; cc: n = 9; vhc+cc: n = 9.

**Supplementary Figure 4.**
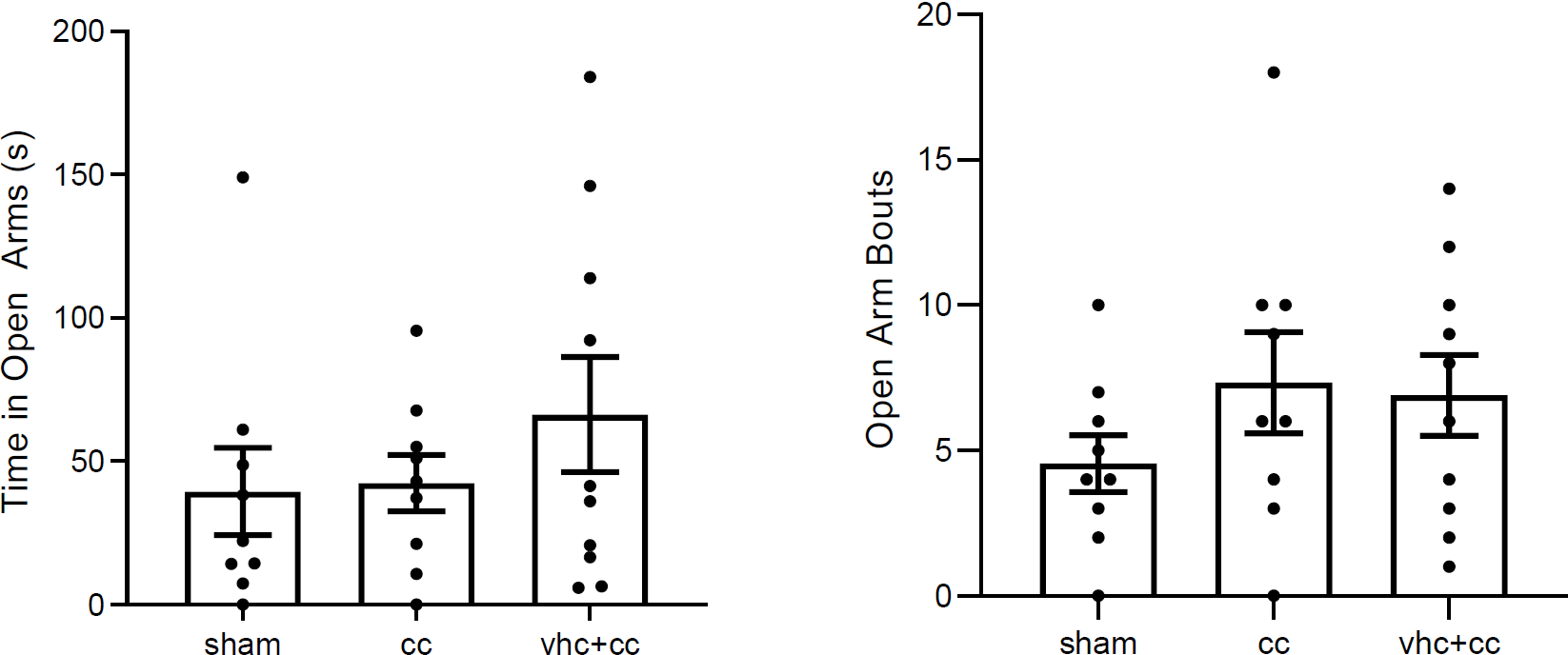
Exploration of the open arms on an elevated plus maze apparatus. There was no difference among groups in time spent in open arms suggesting surgical treatment did not alter anxiety as measured by time spent in the open arms (F(2,25) = 0.867, *p* = 0.433) or the number of open arm bouts (F(2,25) = 1.100, *p* = 0.348). Sham: n = 10; cc: n = 9; vhc+cc: n = 9.

## References

1) Hebb DO. The Organization of Behavior: A Neuropsychological Theory. Oxford, England: Wiley; 1949.

2) Bliss TVP, Collingridge GL. A synaptic model of memory: long-term potentiation in the hippocampus. Nature. 1993; 361: 31–39. DOI: 10.1038/361031a0

3) Tonegawa S, Morrissey MD, Kitamura T. The role of engram cells in the systems consolidation of memory. Nat Rev Neurosci. 2018; 19(8):485–498. DOI: 10.1038/s41583-018-0031-2

4) Turrigiano G. The self-tuning neuron: synaptic scaling of excitatory synapses. Cell. 2008; 135(3):422–35. DOI: 10.1016/j.cell.2008.10.008

5) Bliss TVP, Lømo T. Long-lasting potentiation of synaptic transmission in the dentate area of the anaesthetized rabbit following stimulation of the performant path. J Physiol. 1973; 232: 331–356. DOI: 10.1113/jphysiol.1973.sp010273

6) Kohl, M, Shipton, OA, Deacon, RM, Rawlins, JNP, Deisseroth, K, Paulsen, O. Hemisphere-specific optogenetic stimulation reveals left-right asymmetry of hippocampal plasticity. Nat Neurosci. 2011; 14: 1413–1415. DOI: 10.1038/nn.2915

7) Shipton OA, El-Gaby M, Apergis-Schoute J, Deisseroth K, Bannerman DM, Paulsen O, Kohl MM. Left-right dissociation of hippocampal memory processes in mice. Proc Natl Acad Sci U S A. 2014; 111: 15238–15243. DOI: 10.1073/pnas.1405648111

8) El-Gaby, M, Shipton, OA, Paulsen, O. Synaptic plasticity and memory: New insights from hippocampal left–right asymmetries. Neuroscientist. 2015; 21: 490–502. DOI: 10.1177/1073858414550658

9) Jordan JT, Shanley MR, Pytte CL. Behavior state-dependent lateralization of dorsal dentate gyrus c-Fos expression in mice. Neuronal Signaling. 2020; 3: NS20180206. DOI: 10.1042/NS20180206

10) Jordan JT. The rodent hippocampus as a bilateral structure: A review of hemispheric lateralization. Hippocampus. 2020; 30: 278–292. DOI: 10.1002/hipo.23188

11) Klur, S, Muller, C, Pereira de Vasconcelos, A, Ballard, T, Lopez, J, Galani, R et al. Hippocampal-dependent spatial memory functions might be lateralized in rats: An approach combining gene expression profiling and reversible inactivation. Hippocampus. 2009; 19: 800–816. DOI: 10.1002/hipo.20562

12) Wiltgen BJ, Sanders MJ, Anagnostaras SG, Sage JR, Fanselow MS. Context fear learning in the absence of the hippocampus. J Neurosci. 2006; 26(20): 5484–5491. DOI: 10.1523/JNEUROSCI.2685-05.2006

13) Drew, M. R., Denny, C. A., Hen, R. Arrest of adult hippocampal neurogenesis in mice impairs single-but not multiple-trial contextual fear conditioning. Behav Neurosci. 2010; 124: 446–454. DOI: 10.1037/a0020081

14) Denny CA, Burghardt NS, Schacter DM, Hen R, Drew MR. 4-to 6-Week-old adult-born hippocampal neurons influence novelty-evoked exploration and contextual fear conditioning. Hippocampus. 2012; 22: 1188–1201. DOI: 10.1002/hipo.20964

15) Danielson NB, Kaifosh P, Zaremba JD, Lovett-Barron M, Tsai J, Denny CA, et al. Distinct contribution of adult-born hippocampal granule cells to context encoding. Neuron. 2016; 90: 101–112. DOI: 10.1016/j.neuron.2016.02.019

16) Dillon GM, Qu X, Marcus JN, Dodart J-C. Excitotoxic lesions restricted to the dorsal CA1 field of the hippocampus impair spatial memory and extinction learning in C57BL/6 mice. Neurobiol Learn Mem. 2008; 90(2): 426–433. DOI: 10.1016/j.nlm.2008.05.008

17) Fenton AA, Bures J. Place navigation in rats with unilateral tetrodotoxin inactivation of the dorsal hippocampus: Place but not procedural learning can be lateralized to one hippocampus. Behav Neurosci. 1993; 107: 552–564. DOI: 10.1037//0735-7044.107.4.552

18) Shinohara Y, Hosoya A, Yamasaki N, Ahmed H, Hattori S, Eguchi M et al. Right hemispheric dominance of spatial memory in split-brain mice. Hippocampus. 2012; 22: 117–121. DOI: 10.1002/hipo.20886

19) Maren S, Fanselow MS. Electrolytic lesions of the fimbria/fornix, dorsal hippocampus, or entorhinal cortex produce anterograde deficits in contextual fear conditioning in rats. Neurobiol Learn Mem. 1997; 67: 142–149. DOI: 10.1006/nlme.1996.3752

20) Reis DG, Scopinho AA, Guimarães FS, Corrêa FM, Resstel LB. Involvement of the lateral septal area in the expression of fear conditioning to context. Learn Mem. 2009; 17(3): 134–138. DOI: 10.1101/lm.1534710

21) El-Gaby, M, Zhang, Y, Wolf, K, Schwiening, CJ, Paulsen, O, Shipton, OA. Archaerhodopsin selectively and selectively and reversibly silences synaptic transmission though altered pH. Cell Reports. 2016; 16: 2259–2268. DOI: 10.1016/j.celrep.2016.07.057

22) Spiers HJ, Burgess N, Maguire EA, Baxendale SA, Hartley T, Thompson PJ, O’Keefe J. Unilateral temporal lobectomy patients show lateralized topographical and episodic memory deficits in a virtual town. Brain. 2001; 124: 2476 – 2489. DOI: 10.1093/brain/124.12.2476

23) Maguire, EA, Burgess, N, Donnett, JG, Frackowiack, RSJ, Frith, CD, O’Keefe, J. Knowing where and getting there: A human navigational framework. Science. 1998; 280: 921–924. DOI: 10.1126/science.280.5365.921

24) Howard, LR, Javadi, HR, Yu, Y, Mill, RD, Morrison, LC, Knight, R et al. The hippocampus and entorhinal cortex encode path and Euclidean distances to goals during navigation. Curr Biol. 2014; 24: 1331–1330. DOI: 10.1016/j.cub.2014.05.001

## References

25) kheirbek MA, Drew LJ, Burghardt NS, Costantini DO, Tanneholz L, Ahmari SE, Zeng H, Fenton AA, Hen R. Differential control of learning and anxiety along the dorsoventral axis of the dentate gyrus. Neuron. 2013; 77: 955–968. DOI: 10.1016/j.neuron.2012.12.038

26) Zaidel DW. The case for a relationship between human memory, hippocampus, and corpus callosum. Biol. Res. 1995; 28: 51–57. PMID: 8728820

27) Schalomon PM, Wahlsten D. A precision surgical approach for complete or partial callosotomy in the mouse. Physiol. Behav. 1995; 57: 1199–1203. DOI: 10.1016/0031-9384(94)00349-a

28) Vorhees CV, Williams MT. Morris water maze: procedures for assessing spatial and related forms of learning and memory. Nat. Protoc. 2006; 1: 848–858. DOI: 10.1038/nprot.2006.116

29) Schafe GE, Nadel NV, Sullivan GM, Harris A, LeDoux JE. Memory consolidation for contextual and auditory fear conditioning is dependent on protein synthesis, PKA, and MAP kinase. Learn Mem. 1999; 6: 97–110. PMID: 10327235

30) Kjelstrup KG, Tuvnes FA, Steffenach HA, Murison R, Moser EI, Moser MB. Reduced fear expression after lesions of the ventral hippocampus. Proc. Natl. Acad. Sci. U. S. A. 2002; 99: 10825–10830. DOI: 10.1073/pnas.152112399

